# Diverse cell death signature based subtypes predict the prognosis and immune characteristics within glioma

**DOI:** 10.1101/2024.02.29.582704

**Authors:** Lin Wang, Jia Song, Jing Xu, Yidan Qin, Jia Li, Yajuan Sun, Hui Jin, Jiajun Chen, Ziqian Wang

## Abstract

**Background:** Cell death plays an essential role in the pathogenesis, progression, drug resistance and recurrence of glioma. Although multiple cell death pathways are involved in glioma development, there is lack of a stratification and prognostic modelling for glioma based on the integration of diverse genes for cell deaths.

**Methods:** In this study, 1254 diverse cell death (DCD)-related genes were assessed using the ConsensusClusterPlus assessment to identify DCD patterns in glioma. CIBERSORT, ssGSEA, and ESTIMATE algorithms were applied to evaluate immune microenvironment differences between subtypes. LASSO Cox regression was used to screen prognosis-related DCD genes, and a risk score model was constructed. TMB, TIDE, immune infiltration, and immunotherapy response was analyzed to evaluate the immune characteristics.

**Results:** Two DCD-related subgroups named Clusters 1 and 2, with distinct DCD levels, immune characteristics, and prognoses, were determined from glioma samples. A DCD-based risk score model was developed to assess DCD levels in glioma patients and divide patients into high- and low-risk groups. We found this risk model can be used as an independent prognostic factor for glioma patients. Notably, glioma patients with low risk scores exhibited subdued DCD activity, prolonged survival, and a favorable disposition towards benefiting from immune checkpoint blockade therapies.

**Conclusions:** This study established a novel signature classification and a risk model by comprehensively analyzing patterns of various DCDs to stratify glioma patients and to predict the prognosis and immune characteristics of glioma. We provided a theoretical basis for the clinical application of DCD-related genes in glioma prognosis and immunotherapy.

## Introduction

Gliomas represent a heterogeneous collection of aggressive brain tumors with limited therapeutic options, accounting for most primary central nervous system (CNS) tumors [1]. Their heterogeneity makes it challenging not only for treatment but also for prognosis and immunotherapy-response predictions. In recent years, extensive molecular signatures have been introduced into glioma classification, such as IDH mutation and 1p19q codeletion [2, 3]. Despite these advancement, the vast heterogeneity of gliomas continues to obscure a comprehensive understanding of their classification. Hence, there is an urgent need to identify more specific and practical molecular markers to redefine glioma subtypes and predict the prognosis and response to immunotherapies in glioma patients.

Cell death is a critical event associated with malignant transformation and tumor metastasis [4]. Numerous cell death pathways with different triggering mechanisms and functions have been discovered. Based on functional differences, cell death can be broadly categorized into accidental cell death (ACD) and regulatory cell death (RCD), the latter of which encompasses processes mediated by specific signaling cascades and results in unique biochemical, morphological and immunological events [5]. The RCD that occurs under physiological conditions is called programmed cell death (PCD) or called DCD in this study. PCD/DCD is a kind of intentionally induced cell death accompanied by a series of controlled reactions resulting in programmed self-elimination. Currently known PCD/DCD pathways include apoptosis, necroptosis, pyroptosis, ferroptosis, cuproptosis, parthanatos, entotic cell death, netotic cell death, lysosome-dependent cell death, autophagy, alkaliptosis, and oxeiptosis [6]. Dysregulation of PCD/DCD is commonly associated with the features of malignant tumors, including mortality, metastasis, treatment resistance, recurrence, and tumor immunity [7]. Apoptosis is an intrinsic mechanism of cell death that involves programmed dismantling of cellular components with no impact on the surrounding living cells. Apoptosis is a common tumor suppressor mechanism, whereas apoptosis resistance and malignant proliferation are considered to be important mechanisms in glioma genesis and development [8]. Necroptosis, a form of programmed inflammatory cell death, has recently been confirmed to play a vital role in modulating tumorigenesis and tumor progression [9]. The necroptotic pathway is characterized by RIPK1-RIPK3-MLKL activation downstream of TNFR/Fas and TLR-3/4 [10]. Pyroptosis, another inflammatory PCD pathway induced by some inflammasomes, has been associated with various cancers [11]. Ferroptosis, distinguished by its dependence on iron and extramitochondrial lipid peroxidation, is emerging as a critical modulator of tumorigenesis [12]. Our previous research revealed that cuproptosis-related lncRNAs had an excellent predictive ability for the prognosis and immuno-microenvironment status of glioma patients [13]. Despite these advances, a comprehensive exploration of the relationship between DCD and glioma prognosis remains inadequate, as the detailed roles of DCD in glioma needs further research.

In this study, we aimed to establish a new signature classification and a risk model of DCD-related genes to stratify gliomas and predict the prognosis and immune characteristics of glioma. A series of bioinformatics analyses based on TCGA and CGGA were performed. Based on the expression value of differentially expressed DCD genes, ConsensusClusterPlus was used for a cell death molecular subtype analysis. The CIBERSORT, ssGSEA, and ESTIMATE algorithms were used to evaluate the differences in the immune microenvironment between subtypes. The differences in HLA family genes and immune checkpoint genes between DCD subtypes were analyzed by the Wilcox test. LASSO Cox regression was used to screen prognosis-related DCD genes, and a risk score model was constructed to divide glioma samples into high-risk and low-risk groups. Independent prognostic factor analysis and combined analysis were performed separately to determine whether the risk model could be an independent prognostic factor for glioma. TMB, TIDE, and immunoinfiltration were analyzed to evaluate the immune microenvironment. The pRRophetic algorithm was used to predict the IC50 value for sensitivity prediction analysis of multiple drugs.

In summary, our study identified potential biomarkers reflecting DCD subtype characteristics, which hold potential to become novel prognostic markers and immunotherapy-response predictors for glioma.

## Materials and Methods

### Data collection

RNA-seq data (log2(tpm+0.001)) and clinical information of the corresponding sample were collected from the TCGA-GBM/LGG and GTEx datasets and downloaded from the UCSC-Xena platform. The downloaded clinical information included age, sex, grade, and survival information (OS and OS time). A total of 896 samples (207 paracancerous samples and 689 cancer samples) were included, among which 683 cancer samples had prognostic information and were used as the training set for follow-up analysis. RNA-seq data and corresponding clinical prognostic information from the CGGA database (Chinese Glioma Genome Atlas, http://www.cgga.org.cn/) were downloaded. Samples with OS prognostic information, including 313 glioma tissue samples, were mainly used for verification of the prognostic model.

### Differential expression analysis of DCD-related genes

The DCD-related genes of twelve DCD patterns were collected from GSEA gene sets, KEGG, literature, and manual collation. A total of 1254 genes related to DCD were included in the analysis. After matching with the gene expression matrix of the training set, the expression values of DCD-related genes in 207 paracancer samples and 689 cancer samples were obtained. A differential expression analysis of all DCD-related genes between tumor and normal tissues was performed through linear regression and the empirical Bayesian method provided by the limma package to obtain the corresponding P values. Value and logFC of genes. In addition, the Benjamini & Hochberg method was adopted for multiple inspection and correction, and the p value after correction was obtained, namely, adj.P.Value. The differential expression analysis was assessed according to fold change and significance. Differentially expressed threshold settings were as follows: adj.P. Value < 0.05 & | logFC | > 2. A volcano plot was mapped to display these differentially expressed genes. The R package “clusterProfiler” was used to conduct GO and KEGG enrichment analyses based on the differentially expressed DCD genes.

### DCD Subtype Classification

Based on the expression values of differential DCD genes in each glioma sample obtained above, DCD subtype classification was performed using ConsensusClusterPlus 1.54.0. Then, based on the expression values of all differential DCD genes in the training set, the ssGSEA algorithm was used to calculate the enrichment scores by using the R package “GSVA”. The Wilcox test was used to calculate the significance of the p value in enrichment scores between subtypes, and a violin diagram was drawn.

### Prognosis and clinical correlation analysis of subtypes

Based on the cell death subtypes combined with prognostic information, a Kaplan‒Meier survival curve was produced, and the significance of p value was calculated by the log-rank test. Clinical phenotype information was sorted out to compare clinical information between subtypes. For each factor type variable, a chi-square test was performed to conduct a statistical significance test.

### Immune microenvironment analysis

The immunoinfiltration in tumors is closely related to clinical outcome and has been used as drug targets to improve patient survival. CIBERSORT algorithm was used to evaluate immune microenvironment differences among subtypes. The proportions of 22 kinds of immune cells were calculated based on gene expression levels. The infiltration levels of 28 kinds of immune cells were calculated by using the ssGSEA algorithm and R package “GSVA”. Wilcox test was used to calculate the p values among subtypes, and ESTIMATE algorithm was used to estimate the stromal score, immune score, and ESTIMATE score according to the expression data, and the differences between subtypes were calculated by the intergroup Wilcoxon test.

### HLA family and immune checkpoint gene difference analysis

Expression data of the HLA family and immune checkpoint genes in each tumor sample were extracted. Wilcoxon test was performed to analyze the expression differences of the HLA family and immune checkpoint genes among subtypes.

### GSEA enrichment analysis

First, h.all.v7.4.symbols in the database MSigDB v7.1 were taken as the enrichment background. Based on the gene expression values of each sample and the corresponding subtype information, GSEA was carried out using the R package “clusterProfiler” to determine subtypes that hallmark gene sets significantly enriched in.

### PPI network analysis

The STRING database was utilized to predict interactions among proteins encoded by cell death genes. The input set included differentially expressed cell death genes. The PPI score was set as 0.4 (medium confidence), and the PPI network was constructed using Cytoscape software.

Six topological algorithms (MCC, MNC, Degree, EPC, Closeness, Betweenness) in the cytoHubba plugin of Cytoscape were used to obtain the ordering of genes. The top 20 genes under each algorithm were included for intersection to select key cell death genes.

### Identification of genes associated with prognostic T-cell depletion

In the training set, based on the key cell death genes obtained above, univariate Cox regression analysis was performed to screen out the cell death genes significantly related to overall survival, and a p value < 0.05 was set as the significance threshold.

### Construction and validation of the DCD-related prognostic signature

LASSO and multivariate cox regression analyses were performed successively for further screening of cell death genes related to prognosis. According to the gene regression prognostic coefficients, the Riskscore model is constructed, and the Riskscore calculation formula is as follows:

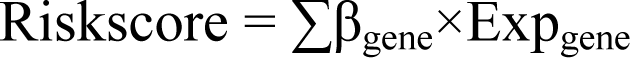

β_gene_ represents the multifactor regression coefficient of each gene, and Exp_gene_ represents the expression level of the gene in the training set.

Furthermore, to verify model accuracy, the Riskscore of each sample in the training set was calculated. According to the median risk score, samples were divided into high- and low-risk groups. The Kaplan‒Meier curve was applied to evaluate the association between the grouping of high- and low-Risk and actual survival. In addition, the ROC curves for 1-, 3- and 5-year prognosis predicted by the RiskScore in the training and validation sets were plotted.

### Correlation of risk scores with clinical characteristics

The distribution box diagram of the risk score in each clinical group (age, sex, grade and cluster) was drawn, and the Wilcoxon test was used to calculate the significance p value between groups (the Kruskal‒Wallis test was used to calculate the significance p value among multiple groups).

### Nomogram establishment

First, to determine whether the Riskscore based model is an independent prognostic factor, univariate cox regression analysis was conducted for age, sex, grade and RiskScore, and variables with a p value < 0.05 significance were included in multivariate cox regression analysis. In the multivariate cox regression analysis, the variables with p values < 0.05 were selected as independent prognostic factors. A nomogram was constructed based on these independent prognostic factors. In addition, calibration and ROC curves were produced to verify the validity of the nomogram.

### Conjoint analysis

According to grade and risk grouping, samples were divided into six groups: high_risk+G2, high_risk+G3, high_risk+G4, low_risk+G2, low_risk+G3, and low_risk+G4. Then, Kaplan‒Meier analysis was used to assess the association between the features and actual prognostic information of the six groups.

### Genomic mutation differences between risk groups

The somatic mutation files of TCGA-GBM and TCGA-LGG datasets were downloaded from TCGA and merged to obtain the TCGA-GBM/LGG somatic mutation maf files. The mutation waterfall maps of the top 20 genes with the highest mutation frequencies in the high- and low-risk groups were plotted using the R package “maftools”. The tumor mutation burden (TMB) of each sample was calculated, and the difference significance between the high- and low-risk groups was calculated by the Wilcoxon test.

### Correlation analysis between diagnostic genes and immunity

Based on the immune infiltration differences between subtypes, the Spearman correlation coefficient and the corresponding significance p. values between model genes and immune cells were calculated, and a correlation heatmap was drawn.

### Predictive analysis of drug sensitivity

Using the pRRophetic algorithm, a ridge regression model was constructed based on the GDSC cell line expression profile and TCGA gene expression profile to predict the IC50 of 138 kinds of drugs. The Wilcox test was used to determine whether there were significant differences in the IC50 of each drug between the high- and low-risk groups.

### Immunotherapy response prediction

Immunotherapy response was predicted by tumor immunodysfunction and elimination (TIDE) analysis. The TIDE tool was used to calculate the TIDE value of each sample in the high-risk and low-risk groups, and the Wilcoxon test was used to calculate the significance of the difference between the groups.

### Immunotherapy data analysis

The GSE91061 dataset was downloaded from the GEO database [20], including 20 samples with a response and 78 samples with no response to immune checkpoint blocking therapy. According to the calculation formula, the Riskscore value of each sample in the dataset was calculated. According to the median risk score, all samples were divided into high-risk (risk score ≥ median risk score) and low-risk (risk score < median risk score) groups. Kaplan‒Meier survival analysis was performed for patients in the High_Risk and Low_Risk groups by using the “survival” package. The Wilcoxon test was used to calculate the significance of the difference in the risk score between the responsive and unresponsive groups.

### Association analysis between cell death subtype and Riskscore

The distribution proportion of cell death subtypes in the High_ and Low_Risk groups was calculated, and significance was calculated by the chi-square test.

## Results

### Differentially expressed gene (DEG) analysis

The workflow of this research is shown in Figure 1. Through the difference analysis and screening threshold setting, 101 differentially expressed genes were obtained, including 86 upregulated and 15 downregulated genes. The volcano map is shown in Figure S1A. In the functional enrichment analysis of differentially expressed DCD genes, we obtained 1917 GO BPs, 123 GO CCs, 165 GO MFs and 59 KEGG pathways. As shown in Figure S1B-C, the top 10 GO and KEGG categories were mainly associated with cell death, inflammatory pathways and the immune response.

**Figure 1.**
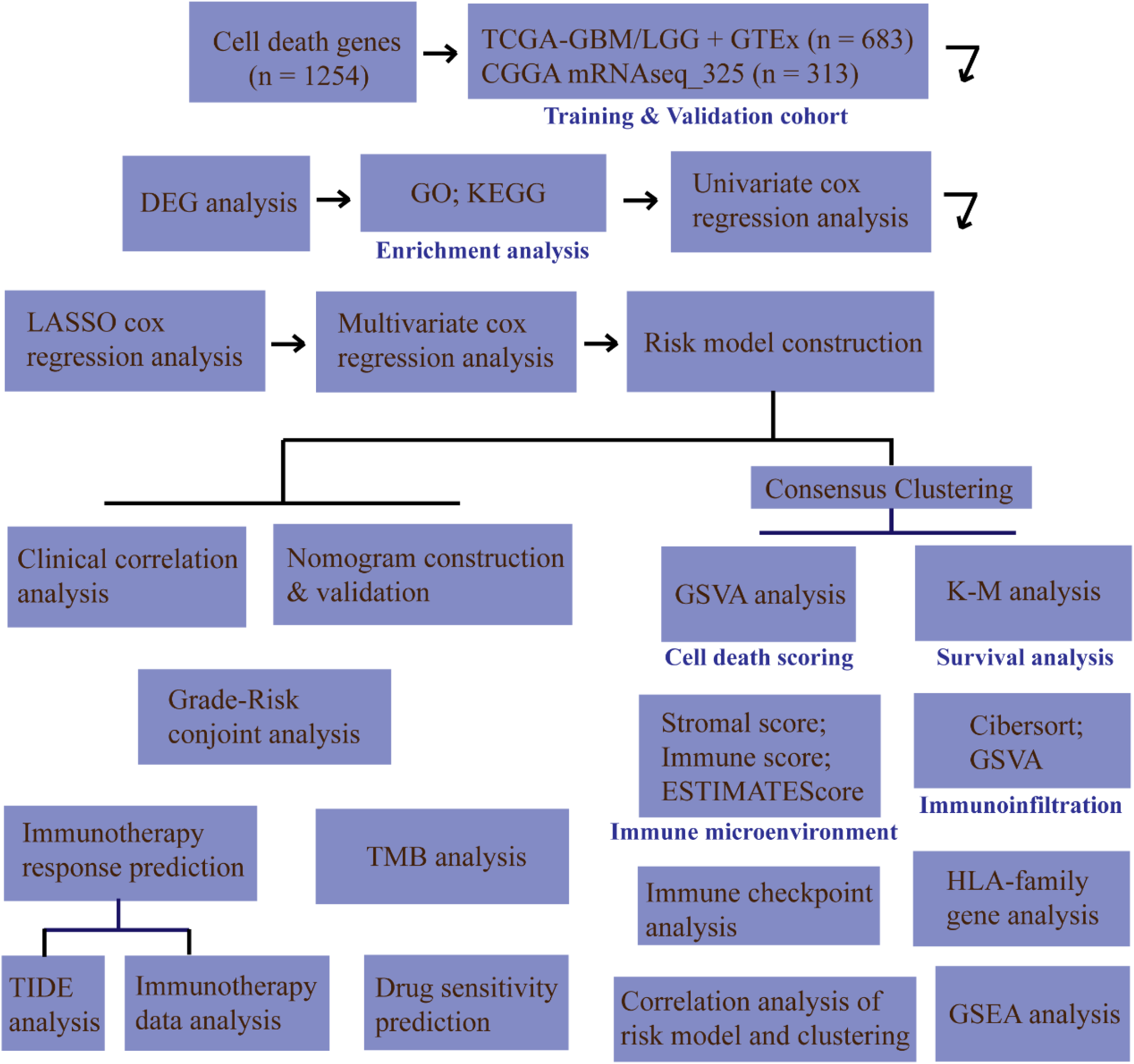
The workflow of this study.

### Identification of DCD-associated molecular subtypes

Based on 101 differentially expressed programmed-cell-death genes (DCDGs) and their expression values in 683 glioma samples with prognostic information, consensus clustering was performed, and two distinguishing clusters were finally determined (Figure 2A-C). The cell death score of each sample was calculated, and combined with the cell death cluster information, we found that the cell death score of Cluster 1 was significantly higher than that of Cluster 2 (Figure 2D). Based on the obtained cell death clusters, the clinical and phenotypic data was sorted out, and the clinical information between clusters was compared, as shown in Table 1, showing that there were significant differences in age and grade between the two clusters. We further investigated the prognosis linked to the two clusters. The overall survival (OS) analysis indicated a significant difference in prognosis between the two clusters, with Cluster 1 having a worse prognosis than Cluster 2 (Figure 2E). Figure 2F shows the differential expression of DCDGs and clinicopathological characteristics between Cluster 1 and 2. Cluster 1 was preferentially associated with higher expression levels of DCDGs and higher grade (G4, Figure 2F).

**Figure 2.**
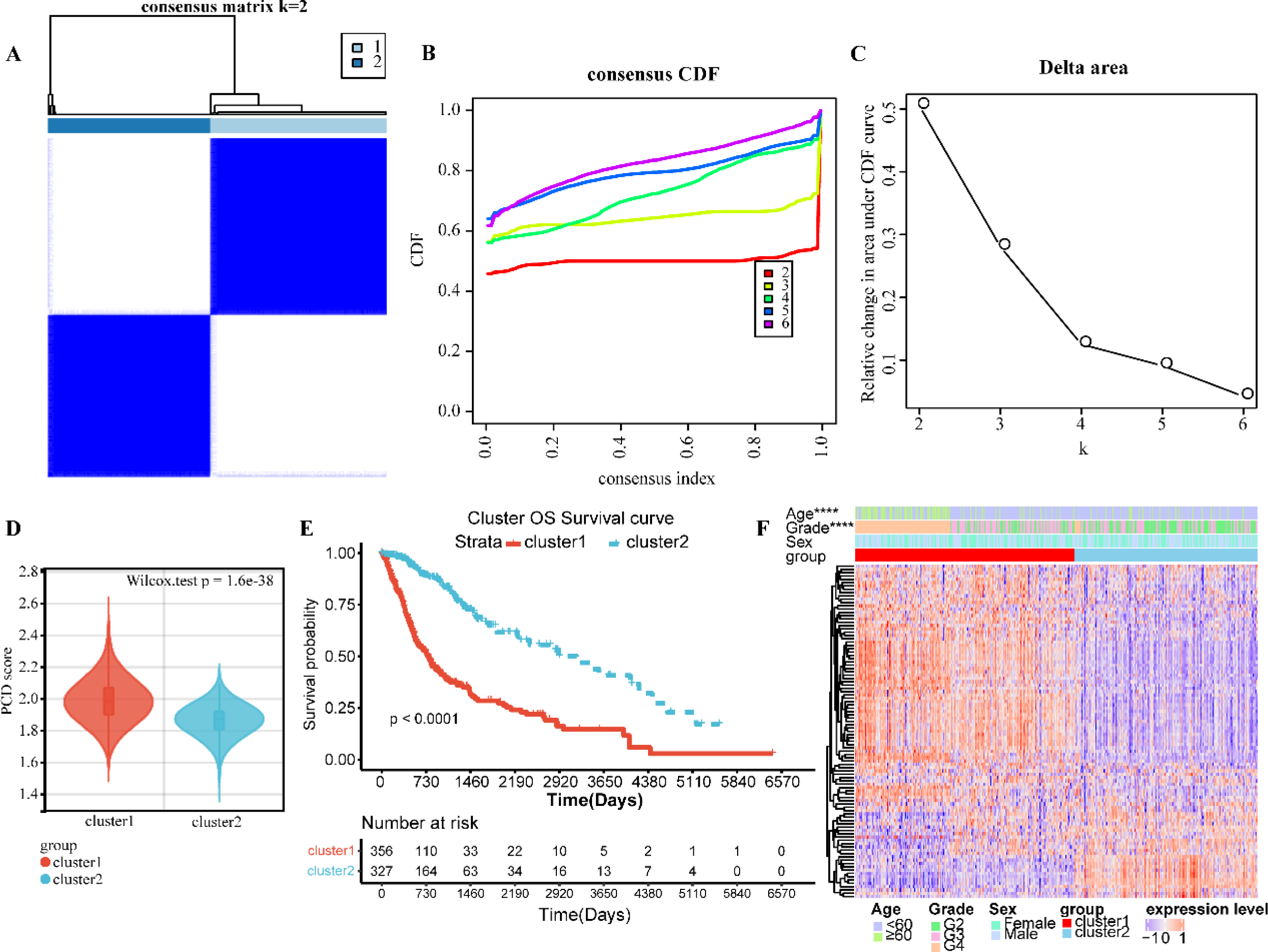
Identification of DCD-associated molecular subtypes. (A-C) consensus matrix heatmap (A), consensus cumulative distribution function plot (B), and delta area plot (C) showed two distinguishing clusters were determined. (D) A violin diagram showed DCD score of Cluster 1 and Cluster 2. (E) The K-M survival analysis of Cluster 1 and 2. (F) A heatmap showed differential expression of DCDGs and clinicopathological characteristics between Cluster 1 and 2.

**Table1.** The clinical information of the two clusters.

| Features | cluster1(n=356) | cluster2(n=327) | pvalue |
| --- | --- | --- | --- |
| Age |  |  | 8.84E-10 |
| <60 | 240 | 286 |  |
| ≥60 | 116 | 41 |  |
| Gender |  |  | 0.4705 |
| Female | 146 | 144 |  |
| Male | 210 | 183 |  |
| Grade |  |  | < 2.2e-16 |
| G2 | 72 | 181 |  |
| G3 | 129 | 135 |  |
| G4 | 155 | 10 |  |

### Immune microenvironment analyses of DCD subtypes

Alternatively, the CIBERSORT algorithm was used to estimate and compare the composition of infiltrated immune cell subpopulations between the two clusters. The analysis showed that monocytes, M2 macrophages, activated dendritic cells, and resting mast cells had a significantly higher and estimated proportion within Cluster 1 than in Cluster 2, while the estimated proportions of naïve B cells, plasma cells, CD4 T cells, and activated mast cells were significantly lower in Cluster 1 than in Cluster 2 (Figure 3A). The ssGSEA algorithm was used to evaluate the infiltration levels of immune cells. We found that the infiltration levels of activated CD4 and CD8 T cells, central memory CD4 and CD8 T cells, macrophages, mast cells, MDSCs, natural killer cells, and regulatory T cells were significantly higher in Cluster 1 than in Cluster 2, while Cluster 2 had higher infiltration levels of monocytes and memory CD4 T cells (Figure 3B). These results revealed different infiltrating immune cell spectra between the two clusters. The high proportion of M2 macrophages and low proportion of M1 macrophages and plasma cells in Cluster 1 suggested that Cluster 1 tends to have an immunosuppressive microenvironment compared with Cluster 2.

**Figure 3.**
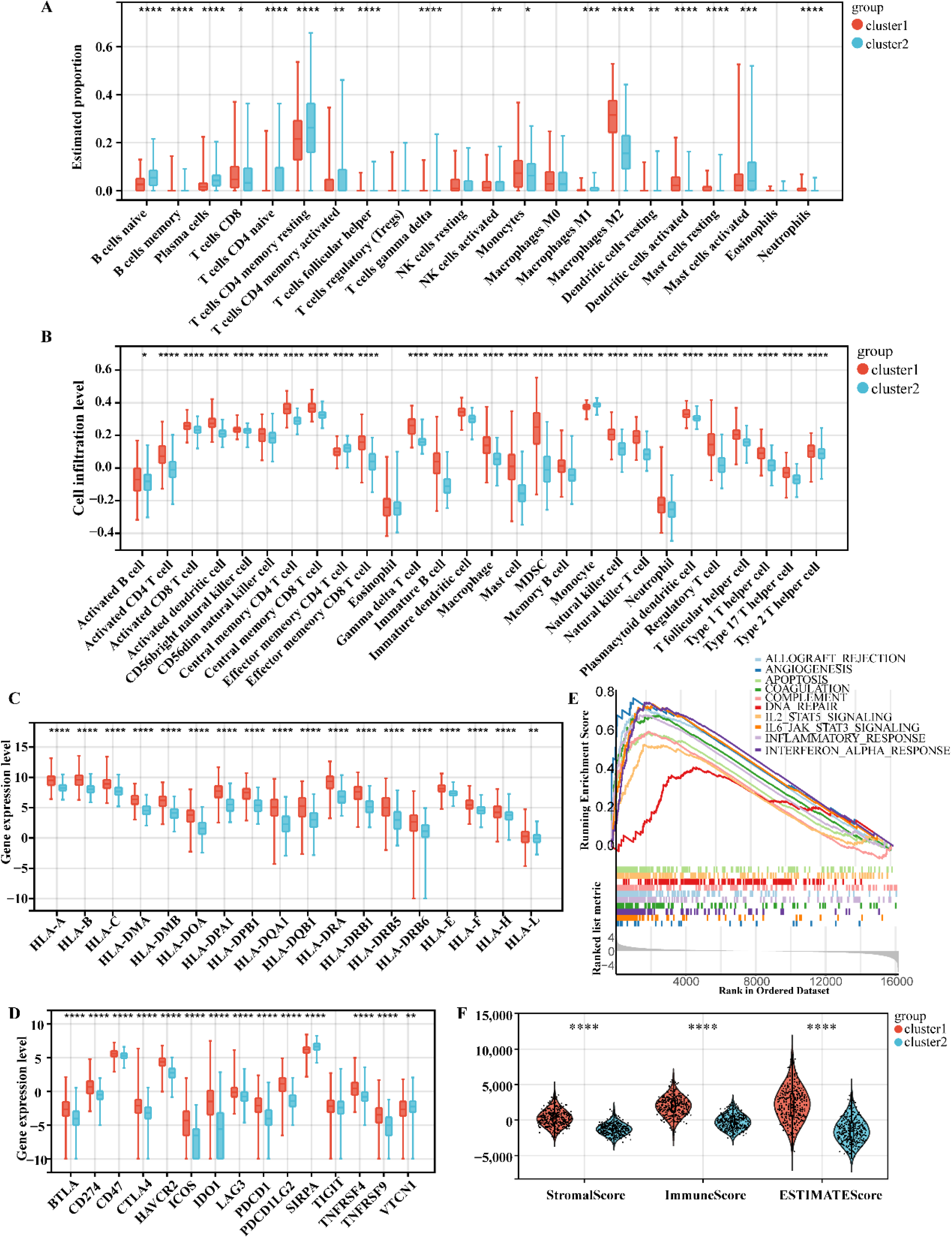
Immune microenvironment analyses of DCD subtypes. (A) Immune cell distribution box diagram based on the CIBERSORT algorithm. (B) Immune cell distribution box diagram based on the ssGSEA algorithm. (C) A distribution box diagram of HLA family genes. (D) A distribution box diagram of immune checkpoint genes. (E) GSEA enrichment results of up-regulated hallmark genes (Cluster1 VS Cluster2). (F) The distrabution violin diagram showing immune score, stromal score and estimate score of two clusters.

Generally, tumors with higher HLA gene expression have more abundant infiltration of immune cells. However, the relationship between the expression of HLA family genes and the immune microenvironment and prognosis of glioma remains controversial. Here, we found significant expression differences in 18 HLA genes between the two clusters. Cluster 2 had lower expression levels of HLA genes than Cluster 1 (Figure 3C). Among them, the HLA-DR expression level is closely related to the invasiveness of glioma and is positively correlated with the glioma grade. In addition, high levels of HLA-F expression are thought to be associated with higher grade and poorer prognosis of glioma. The gene expression levels of immune checkpoint genes (ICGs), including BTLA, CD274, CD47, CTLA4, HAVCR2, ICOS, IDO1, LAG3, PDCD1, PDCD1LG2, TNFRSF4, and TNFRSF9, in Cluster 1 were significantly higher than those in Cluster 2 (Figure 3D), while SIRPA and VTCN1 had lower expression levels in Cluster 1 than in Cluster 2. The above results suggested an immune-suppressive and tumor-supportive microenvironment of Cluster 1. Then, hallmark gene enrichment of Cluster 1 *vs.* 2 was analyzed, and a total of 30 upregulated and 2 downregulated hallmark gene sets were enriched. The top 10 upregulated hallmark gene sets are shown in Figure 3E, including gene sets related to the interferon-α response, inflammatory response, IL-2_STAT5 signaling, IL-6_JAK_STAT3 signaling, allograft rejection, complement, and so on. Additionally, the ESTIMATE algorithm was used to estimate the immune score, stromal score, and microenvironment score. As shown in Figure 3F, the immune score, matrix score and microenvironment score were all significantly higher in Cluster 1 than in Cluster 2, indicating that Cluster 1 may have a more abundant infiltration of nontumor cells than Cluster 2.

### Differentially expressed gene analyses of DCDGs

PPI analysis was performed on 101 differentially expressed DCDGs, including 385 PPI pairs and 95 gene protein nodes (Figure S2A). Six topological algorithms (MCC, MNC, degree, EPC, closeness, betweenness) under Cytoscape were used for the analyses to obtain the ordering of genes. The top 20 genes under each algorithm were selected for intersection, and 10 key DCDGs were finally screened out, as shown in Figure S2B-G.

### Identification of prognostic DCDGs

Based on the 10 key DCDGs obtained above, univariate cox regression analysis was conducted, and the results showed that all 10 DCDGs had p values less than 0.05 (Figure 4A). Furthermore, combined with the expression values of the 10 genes in glioma samples as well as the survival time and survival state of the samples, 9 optimized feature genes were screened by LASSO cox regression, as shown in Figure 4B-C. Then, stepwise multifactor cox regression analysis was performed on the 9 characteristic genes, and 8 key genes related to prognosis were obtained and are shown in the forest plot, including CD44, CD68, CD74, CYBB, IL-18, IL1B, SYK, and TP53 (Figure 4D).

**Figure 4.**
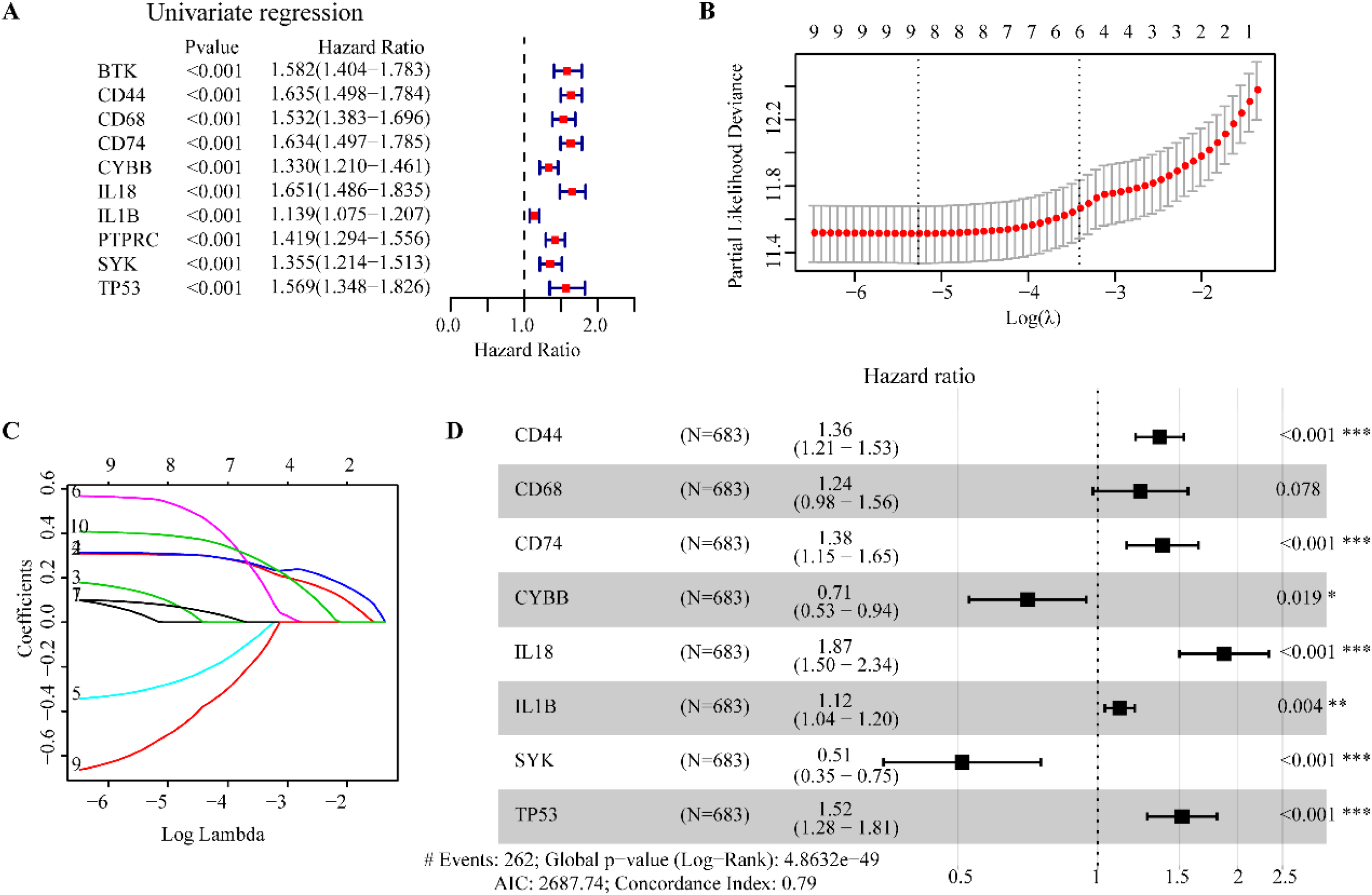
Identification of prognostic DCDGs. (A) The univariate Cox regression forest plot of 10 DCDGs with significant prognostic correlation. (B-C) Lasso regression analysis screened 9 DCDGs. (D) Multivariate COX regression analysis identified 8 DCDGs related to prognosis.

Based on the multivariate regression coefficients of the 8 genes and their expression levels, a risk model was constructed to obtain the risk scores of the samples in the training set and CGGA mRNAseq_325 verification set. Risk score visualization is shown in Figure 5A-B. According to the median Riskscore value, the samples were divided into high- and low-Risk groups. Kaplan‒Meier survival analysis was conducted to evaluate the association between grouping and actual prognosis, as shown in Figure 5C-D. It is obvious that the high-risk patients have shorter survival times than the low-risk patients. Furthermore, 1-, 3- and 5-year survival ROC curves were produced according to survival information and risk scores (Figure 5E-F).

**Figure 5.**
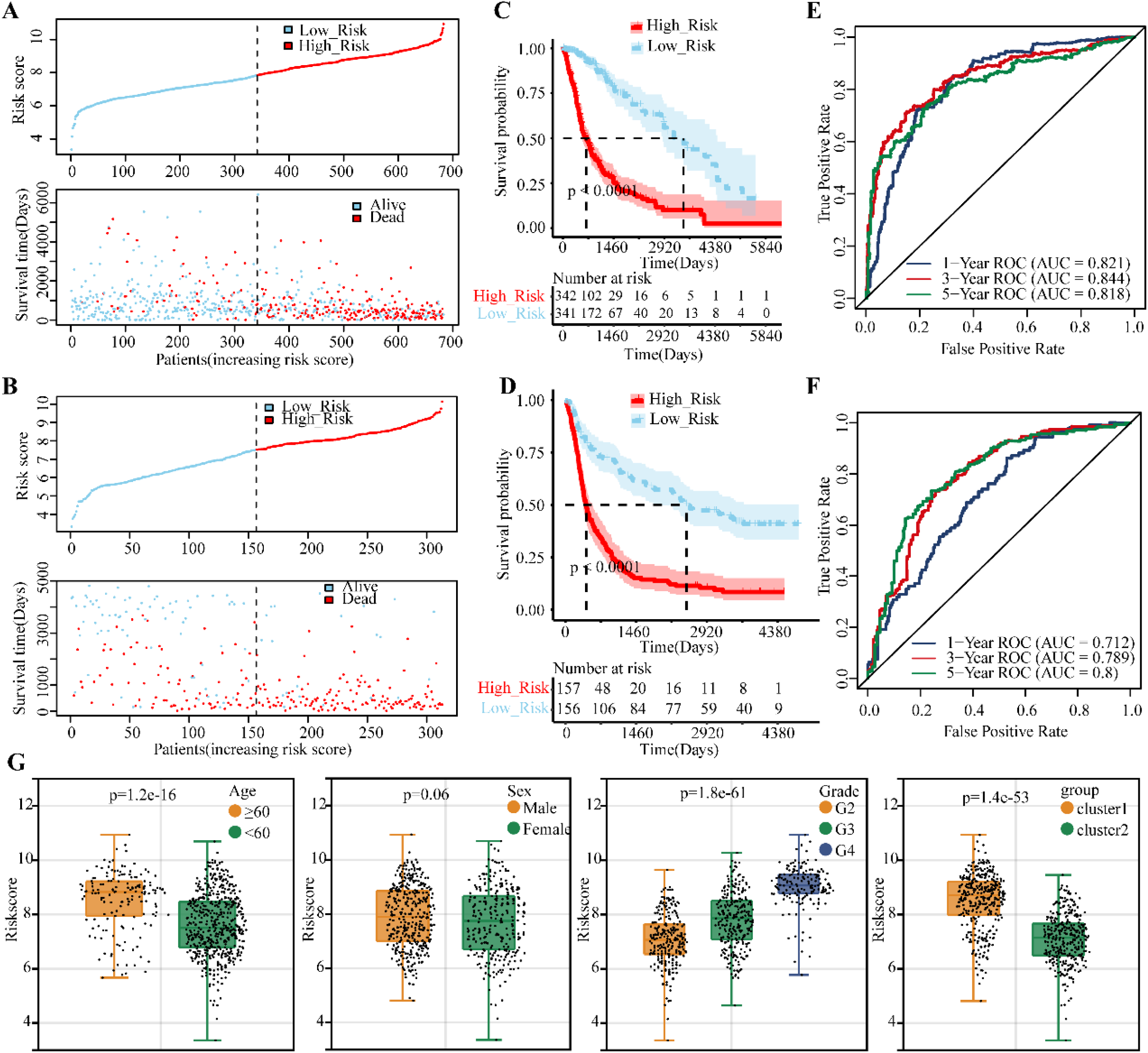
Evaluation of the prognostic significance of risk score. (A-B) For the training set (A) and CGGA mRNAseq_325 verification set (B), sample plots with risk score ranked from low to high and the corresponding survival time of each sample. (C-D) Survival curve of the training set (C) and CGGA mRNAseq_325 verification set (D). (E-F) ROC curves of the training set (E) and CGGA mRNAseq_325 verification set (F). (G) Distribution box plots of the risk score under different clinical factors.

The AUC values are 0.712, 0.789, and 0.8 for 1-, 3-, and 5-year survival in the verification set. The results showed that there was a significant correlation between grouping by risk score and actual prognosis. In addition, the risk score box plots under each clinical characteristic group (age, sex, grade) and among clusters were drawn and are shown in Figure 5G. This result indicated that there were significant differences in the risk scores of different clusters, grades, or ages.

Next, univariate cox regression analysis was carried out for age, sex, WHO grade, IDH status, MGMT status, and risk score (Figure 6A). Variables with p values < 0.05 were included in multivariate cox regression analysis, and then factors with p values < 0.05 were further selected (Figure 6B). The nomogram in Figure 6C showed that age, grade, IDH status, and RiskScore were independent risk indicators for gliomas. The calibration curve and ROC curve proved the accuracy of the nomogram (Figure 6D-E).

**Figure 6.**
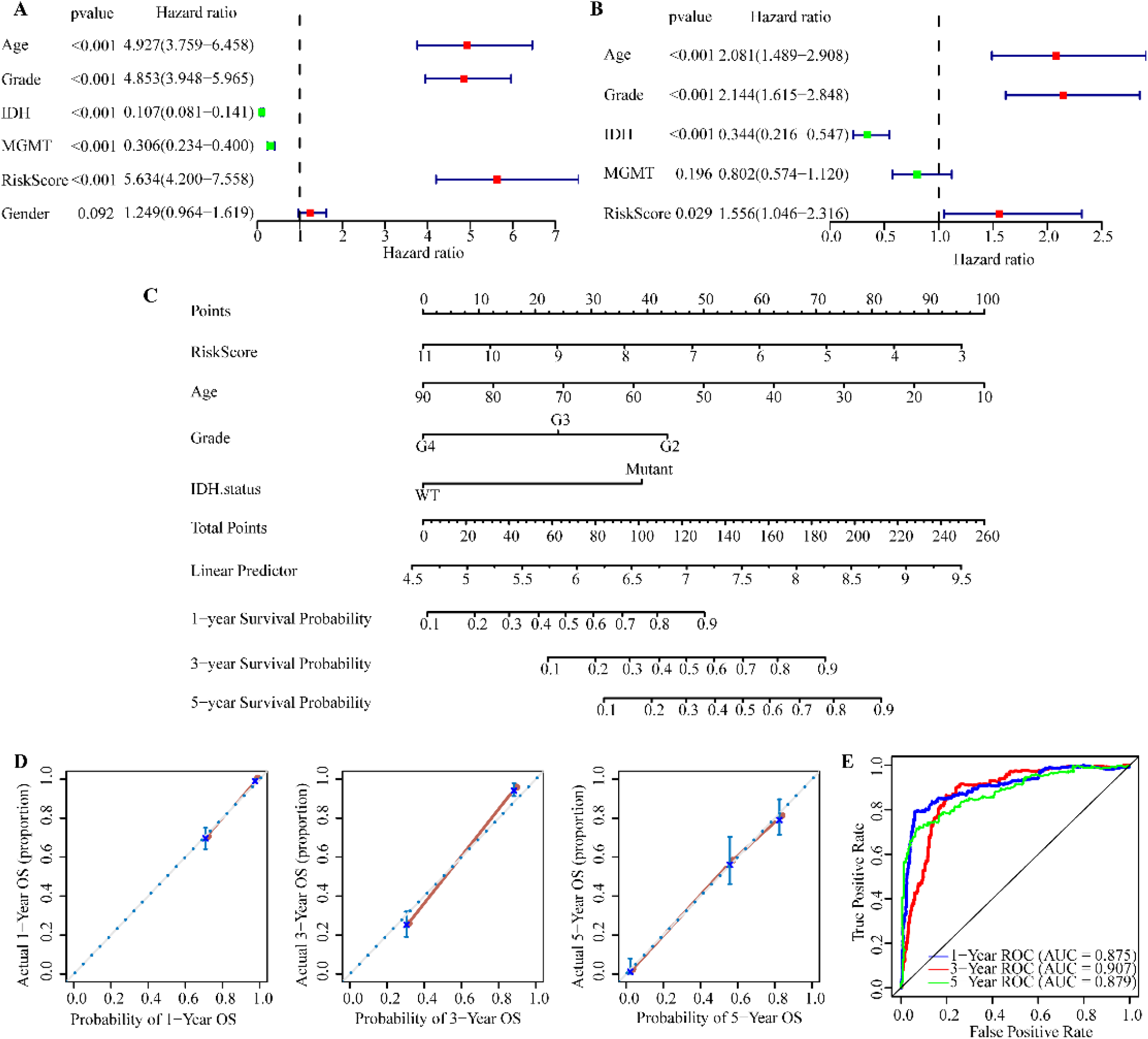
Analysis of independent prognostic factors. (A-B) Forest plots showing the univariate (A) and multivariate (B) regression analyses of clinical factors. (C) The normgram showing independent risk indicators for gliomas. (D-E) The normgram calibration curve (D) and ROC curve (E).

### TMB and drug sensitivity prediction analyses

According to the risk grouping and glioma grade, the samples were divided into 6 groups, and K‒M survival analysis was conducted and is shown in Figure 7A. The results revealed that both high grade and high risk score was associated with short survival. In each grade, the high-risk group had a poorer prognosis as compared with the low-risk group. The top 20 genes with the highest mutation frequencies were shown in oncoplots in Figure 7B-C, showing that high-frequency mutations were significantly different between the high- and low-risk groups. The tumor mutation burden (TMB) score results showed that the high-risk group had significantly higher TMB score than the low-risk group (Figure 7D). In general, tumors with high TMB tend to be more aggressive and progress faster. In addition, the percentages of the two clusters were calculated and shown in Figure 7E. We found that the high-risk group contained more Cluster 1 samples than the low-risk group. In view of the immunosuppressive and tumor-supporting characteristics of Cluster 1, the results suggested that the risk score matched the status of tumor-induced immunosuppression. Then, a drug sensitivity prediction analysis was performed to calculate the half-maximal inhibitory concentration (IC50). The results indicated that 125 drugs showed significant differences in IC50 values between the high- and low-risk groups. The different IC50 values of the common drugs rapamycin, axitinib and gefitinib were shown in Figure 7F-H.

**Figure 7.**
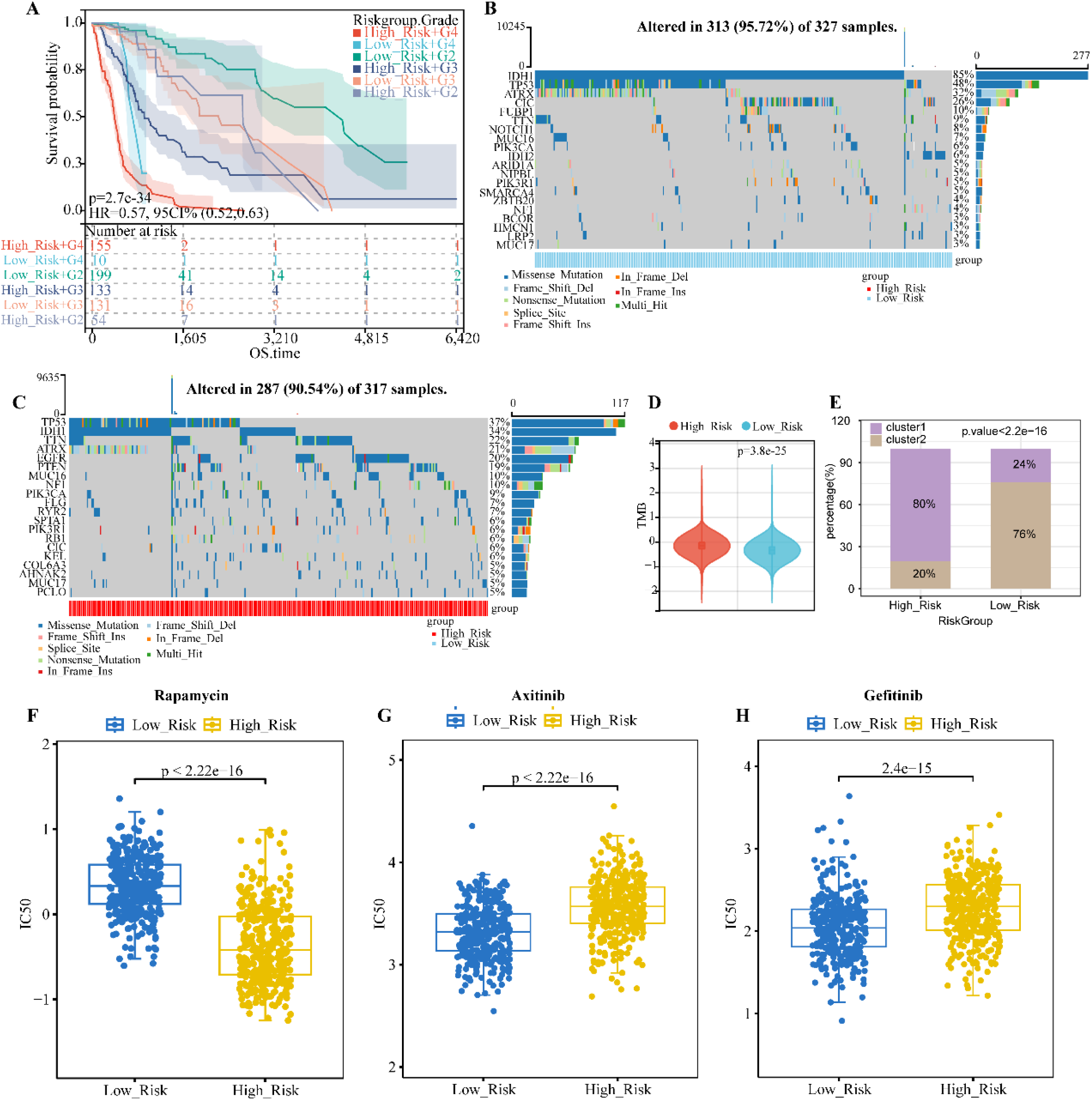
TMB analysis and drug sensitivity prediction analysis. (A) Combined survival analysis of risk and grade for glioma. (B-C) Waterfall plots showing the TOP20 genes with the highest mutation frequencies in the low- (B) and high-risk (C) groups. (D) A violin plot showing the TMB distribution of the low- and high-risk groups. (E) Percentages of the two DCD subtypes in the low- and high-risk groups. (F-H) The box plots showing IC50 values of rapamycin (F), axitinib (G), gefitinib (H) in the low- and high-risk groups.

### Immune microenvironment analyses on the risk model

First, Spearman correlation coefficients between model genes and immunoinfiltrating cells and the corresponding p values were calculated. The correlation heatmaps in Figure 8A-B showed that the model genes were positively correlated with the infiltration of effector-memory-CD8-T cell, neutrophil, type-1-T-helper-cell, natural-killer-cell, macrophage M2, activated-dendritic-cell and so on. TIDE represents tumor immune dysfunction and rejection and is commonly used to assess the likelihood of tumor immune escape. The violin plot in Figure 8C revealed a significant difference in the TIDE score between the high- and low-risk groups. This result suggested that the high-risk group may have a stronger immune escape potential than the low-risk group. The risk score of samples in GSE91061 was calculated, and a risk score distribution box plot of the response and nonresponse groups was produced. The results showed that the nonresponse group had a higher risk score than the response group (Figure 8D). Furthermore, the proportion of response and nonresponse was counted, and we found that the high-risk group contained more nonresponse patients (Figure 8E). Above results indicated that the high-risk group was less responsive to immunotherapy than the low-risk group. In addition, the high-risk group showed poorer prognosis than the low-risk group (Figure 8F). These results revealed that the high-risk group had more significant immune escape characteristics and a lower likelihood of immunotherapy benefit than the low-risk group.

**Figure 8.**
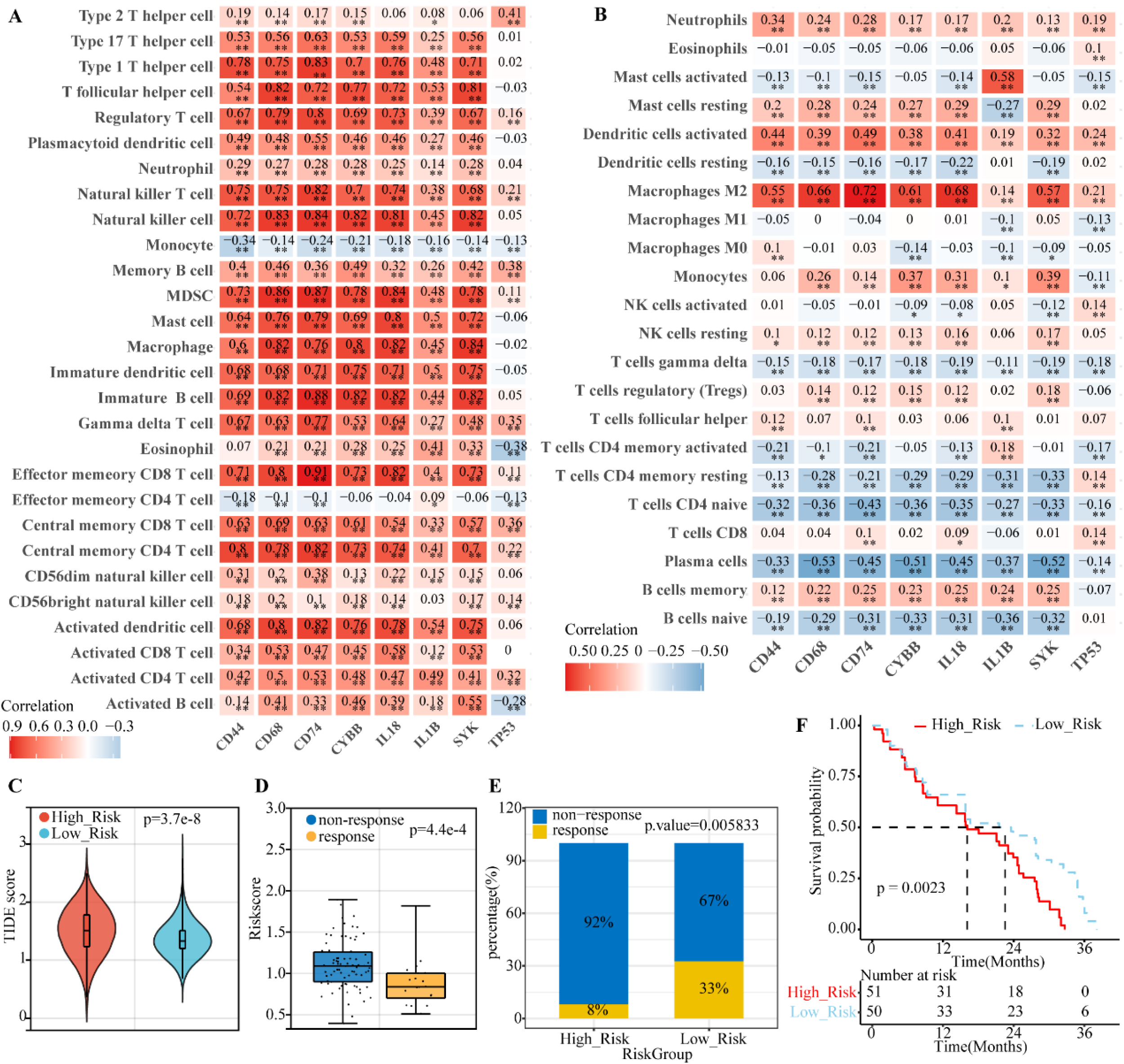
Immune microenvironment analyses on the risk model. (A-B) Heatmaps showing correlations of 8 model genes with infiltrating immune cells using ssGSEA (A) and cibersort algorithm (B). (C) TIDE score of the low- and high-risk groups. (D) The risk score of the non-response and response groups. (E) Percentages of non-response and response in the low- and high-risk groups. (F) K-M survival curve for the low- and high-risk groups in the GSE91061 dataset.

## Discussion

By estimation, gliomas account for approximately 80% of central nervous system malignant tumors worldwide [14]. Despite improvements in diagnosis and treatment in recent years, the prognosis of gliomas, especially glioblastomas, remains unsatisfactory.

Cell death is a necessary condition to maintain the growth and development of organisms [15]. One of the critical characteristics of tumor cells is it’s resistance to cell death. By resisting death and avoiding immune killing, tumor cells achieve continuous division and proliferation out of control of the normal growth regulatory system [4]. However, the hypermetabolism of tumor cells causes a relative shortage of oxygen and nutrients needed for tumor growth and induces necrotizing cell death in the interior of solid tumors [16]. With new discoveries of cell death pathways and related mechanisms, the understanding of the role of cell death in tumors is constantly deepening. The prognostic value of multiple DCD-related genes in malignant tumors, including glioma, has been consistently validated. For example, Wang et al. constructed a risk model based on cuproptosis-related lncRNAs, which showed good prognostic prediction performance and indicated the immuno-microenvironment status for glioma [13]. Chen et al. identified 11 ferroptosis-related genes highly correlated with the prognosis of glioma patients[17]. However, there is a lack of a comprehensive evaluation of all DCD-related genes in terms of their prognostic value, strata performance and immunologic characteristics in glioma, which is the scientific problem our study aims to address.

Numerous studies have demonstrated that infiltrating immune cells in the microenvironment exert dual effects of promoting tumor and antitumor growth [18, 19]. On account of the destruction of blood‒brain barrier integrity by gliomas and the lymphatic outflow channels, the immune system can communicate with cells within the central nervous system [20, 21]. The immune infiltration of gliomas is characterized by extensive spatial and molecular heterogeneity. Gliomas can be infiltrated by a variety of immune cells, including macrophages, microglia, B cells, T cells, myeloid suppressor cells (MDSCs), etc.[22]. Previous studies have shown that the number of microglia and macrophages is positively correlated with glioma grade and invasiveness [23]. Our study revealed that both monocytes and M2 macrophages showed significantly higher infiltrating levels in Cluster 1 than in Cluster 2. Monocytes infiltrate the tumor and differentiate into tumor-associated macrophages and dendritic cells, which influence the tumor microenvironment through various mechanisms and result in immune tolerance, angiogenesis and metastasis [24]. M2 macrophages synthesize and release many anti-inflammatory factors (such as IL-10 and TGF-β).), immunosuppressive factors and a variety of cytokines, inhibiting the inflammatory response and promoting tumor growth and metastasis [25]. In addition, another type of cancer-promoting immune cell, type-2 helper (Th2) cells, was upregulated in Cluster 1 compared with Cluster 2. Alternatively, some antitumor immune cells, such as plasma cells, M1 macrophages and activated mast cells, were upregulated in Cluster 2. The results showing more abundant M2 macrophages and Th2 cells but fewer M1 macrophages and plasma cells in Cluster 1 was consistent with the analysis that Cluster 1 showed a poorer prognosis than Cluster 2. For most HLA genes and immunological checkpoint genes (ICGs), their expression levels in Cluster 1 were higher than in Cluster 2. Therefore, these HLA genes and ICGs are expected to be potentially effective therapeutic targets for Cluster 1.

In most cancers, a higher TMB represents a better response to immune checkpoint suppression therapy. However, high TMB in gliomas has rarely been reported to be associated with better survival outcomes in response to immunotherapy. In a study published in 2021 [26], in two queues layered by TMB, patients with recurrent GBM (rGMB) with a TMB ≤ median lived longer after anti-PD-1/PD-L1 treatment than patients with a TMB > median. Therefore, TMB may not be an independent predictor of the response to immunotherapy in glioma. Instead, we used TIDE to predict the response to immunotherapy. The TIDE score provides a better assessment of the efficacy of anti-PD1 and anti-CTLA4 therapies than widely used biomarkers (TMB, PD-L1, and interferon- γ). In addition, TIDE was stable in predicting efficacy regardless of the tumor-infiltrating level of cytotoxic T cells [27]. Our analyses suggested that high risk tends to be associated with immunotherapy nonresponse compared to low risk. It should be noted that since DCD contains a wide variety of cell death pathways, the distinct DCD levels may cause different effects, leading to dual roles in glioma. When the DCD level is not sufficient to induce cell death, glioma cells may activate intrinsic signaling pathways as response to stimulation and resist death. However, sufficient levels of DCD to induce cancer cell death may be a promising option for glioma therapies. Nevertheless, there are still some limitations. First, phase 3 randomized controlled trials are lacking in this study, so the decision-making role and strata performance of this DCD-related model in a specific patient population cannot be verified. Second, the biological functions of some DCD genes need to be investigated intensively in both in vivo and in vitro experimental studies to fully understand their roles in the pathogenesis and progression of glioma.

In conclusion, we established a DCD-related signature classification model and a DCD-based risk model to predict the prognosis and intratumor microenvironment for glioma patients, which is of significance for the development of glioma treatment strategies.

## Funding Information

This work was supported by the National Natural Science Foundation of China [82203647], Special Project of Health Research Talents of Jilin Province [2022SC234], and Norman Bethune Program of Jilin University [2022B28].

## Conflict of Interest

No potential conflict of interest was reported by any authors of this research.

## Data availability

Data available on request from the authors.

## Notes

### Competing Interest Statement

The authors have declared no competing interest.

